# Conservation motifs - a novel evolutionary-based classification of proteins

**DOI:** 10.1101/2020.01.12.903138

**Authors:** Hodaya Beer, Dana Sherill-Rofe, Irene Unterman, Idit Bloch, Mendel Isseroff, Doron Stupp, Elad Sharon, Elad Zisman, Yuval Tabach

## Abstract

Cross-species protein conservation patterns, as directed by natural selection, are indicative of the interplay between protein function, protein-protein interaction and evolution. Since the beginning of the genomic era, proteins were characterized as either conserved or not conserved. This simple classification became archaic and cursory once data on protein orthologs became available for thousands of species.

To enrich the language used to describe protein conservation patterns, and to understand their biological significance, we classified 20,294 human proteins against 1096 species. Analyses of the conservation patterns of human proteins in different eukaryotic clades yielded extremely variable and rich patterns that had never been characterized or studied before. Using mathematical classifications, we defined seven conservation motifs: *Steps, Critical, Lately Developed, Plateau, Clade Loss, Trait Loss* and *Gain*, which describe the evolution of human proteins.

One type of motif, which we termed *Gain*, describes the human proteins that are highly conserved in a small number of organisms but are not found in most other species. Interestingly, this pattern predicts 73 possible instances of horizontal gene transfer in eukaryotes.

Overall, our work offers novel terms for conservation patterns and defines a new language intended to classify proteins based on evolution, reveal aspects of protein evolution, and improve the understanding of protein functions.

## Introduction

Creating a more accurate terminology and precise nomenclature can facilitate a deeper perspective on biological processes. One of the revolutions in system biology resulted from the definition and study of network motifs^1^. Network motifs are defined by a particular pattern of interactions between proteins that reflect functional properties. A contribution to the protein-protein interaction field was also presented, suggesting two types of networks (hubs): ‘party’ and ‘date’^2^. These networks can better define protein dynamic connectivity in a single organism over time. Based on these definitions, new analyses have become possible, contributing greatly to our understanding of protein interaction and function. In a similar manner, numerous paths exist to characterize and classify proteins, which improved the ability to assign function to poorly characterized proteins. These include classification of proteins into functional annotations^3–5^, subcellular localization and organismal levels^6–8^, and classification of domains and functional sites^9–11^. These characterizations generated a unified language now applied by the scientific community, allowing better communication and resulting in advances the relevant field of research. Although many aspects of protein characterization have been analyzed, no characterization uses information related to protein conservation and evolution. In this work, we offer a new layer to protein classification, based on evolutionary patterns, as revealed by phylogenetic profiling data.

Phylogenetic profiling is a mathematical representation that describes the evolution of a protein as a pattern of its conservation (i.e. presence or absence) in a set of species^12^. A recent approach with a continuous scale for protein conservation is the Normalized Phylogenetic Profiling (NPP)^13,14^. The NPP assigns, for each of the human proteins, a score representing its relative conservation (between 0 to 1) in each species. Over the past decade, NPP has been used to discover genetic interactions and novel genes in the RNA interference pathway, RNA methylation, and various human diseases including cancer^13–20^. Recently we demonstrated the importance of focusing on different taxonomic clades as it can improve the ability to identify novel factors in different pathways, specifically in the homologous recombination repair pathway^21^.

Despite an exponential accumulation of protein sequences from thousands of genomes, protein conservation is still defined essentially by Boolean terms (Fig. 1-A). This definition of conservation is very subjective, as “conserved” or “not conserved” may differ when comparing orthologs of two organisms or of multiple organisms. Furthermore, the comparison may be between closely related organisms (such as human and mouse) or between very distant ones (such as human and E. coli). Moreover, in nature, proteins have very complex evolutionary trajectories such that, within the same evolutionary timeframe, a protein can be fully lost, diverge dramatically, or stay almost without changes throughout the tree of life or in specific branches (clades) (Fig. 1-B). However, no objective criteria exists that includes the above information.

**Figure 1:**
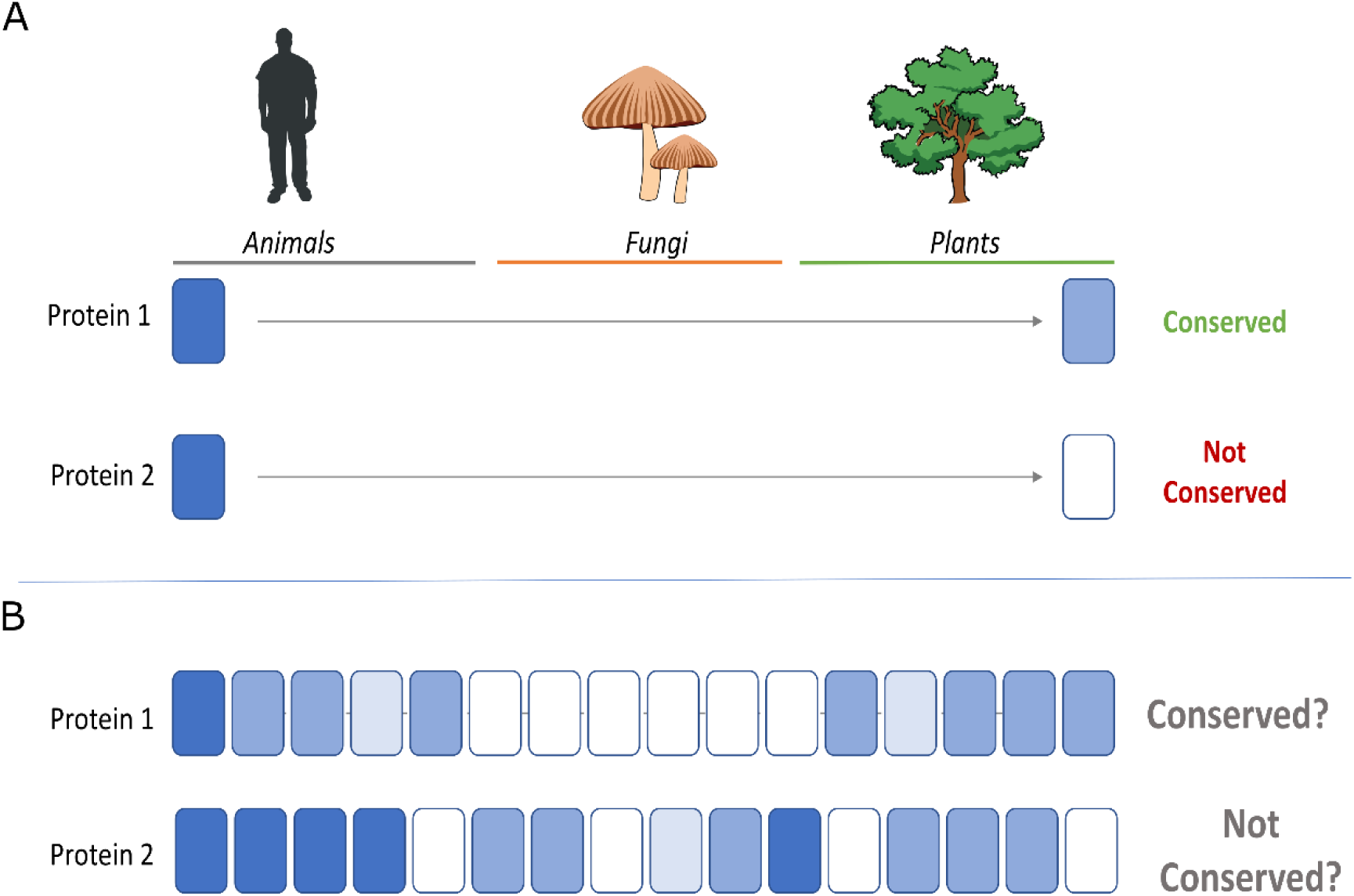
Information hidden in a gene’s evolutionary history. A demonstration of the signal lost by using partial information and simplistic terms for describing protein evolution. Dark blue – high conservation score, light blue – low conservation. (**A**) The illustration presents two human proteins; protein 1 is conserved in a distant organism while protein 2 is not. Although this description seems adequate and correct, it is rather simplistic. (**B**) Representation of the complexity of protein conservation as observed from the LNPP data. When considering the continuous scores among multiple species, a new layer for the term ‘conservation’ can be added.

Phylogenic profiling analysis is commonly used to detect functionally related proteins based on correlated evolution. However, detecting protein conservation motifs can reveal much more than co-evolved proteins as it can shape the overall evolutionary concepts of proteins. Conservation motifs reflect protein function, interaction and selective pressure undergone throughout evolution. An intuitive example for a conservation pattern of a protein may be high conservation among all, or most, species. The evolutionary insight about this *Critical* motif is that the protein serves a crucial role in many species survival and may indicate a role for this protein as a “house-keeping” gene. Other proteins show conservation patterns that correlate with the evolutionary distance of the species. In contrast, some proteins show complex patterns such as massive loss or conservation variability within taxonomic clades.

Here we identify and define recurring conservation motifs of almost all human protein-coding genes across eukaryotes. We show that conservation motifs can define and frequently explain proteins changes along evolution, adding a new layer to traditional protein evolution terminologies. We show that conservation motifs encompass biological insights with regard to the proteins role based on their continuous evolutionary score.

## Results

Our goal was to identify recurring evolutionary patterns, termed conservation motifs, throughout 1096 species in the tree of life. We searched for common conservation patterns, which we formulated using mathematical criteria. The mathematical criteria recapitulate significant information about protein evolution and the effect of selective pressures on proteins. The motifs were defined a-priori in order to capture a spectrum of evolutionary behaviors. The species include 8 taxonomy clades: Primates, Other Mammalians, Other Chordata, Other Metazoa, Alveolata, Fungi, Viridiplantae and Other Eukaryotes^21^. These clades were designed to capture a human-centric understanding of protein evolution.

The conservation motifs can be analyzed using two approaches: **Global**, which considers the full profile of the protein among all, or most, of the species, and **Local**, which considers specific regions of the profile within specific taxonomic groups. The motifs mathematical criteria were formulated to be suitable for a global and a local analysis, according to one’s field of interest. However, some of the motifs are more global and some are more local by definition (Fig. 2).

**Figure 2:**
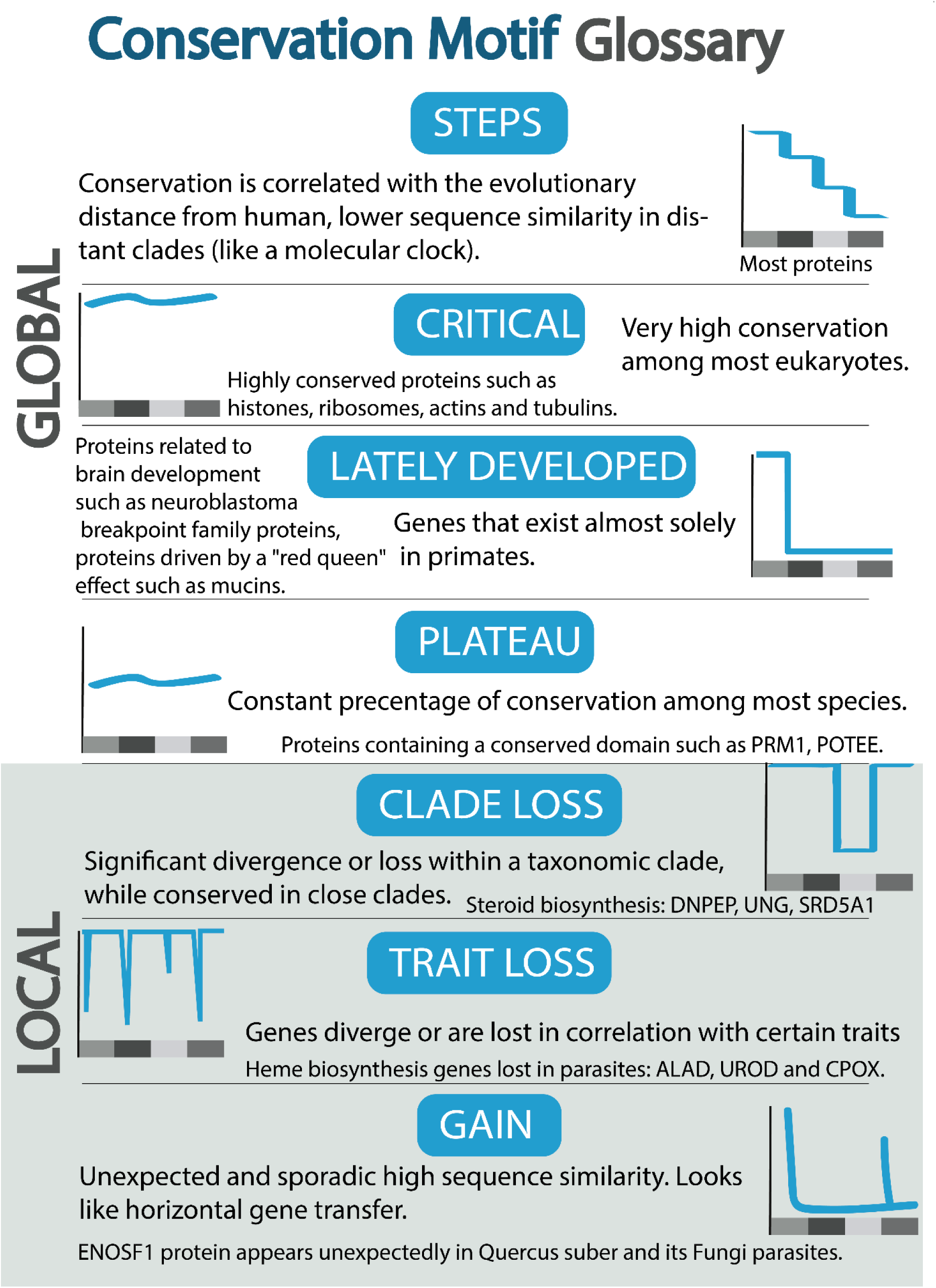
Conservation motifs glossary. The two main categories of motifs are: ‘Global’ and ‘Local’. The global category includes four motifs: Steps, Critical, Lately developed and Plateau. The local category includes three motifs: Clade loss, Trait loss and Gain. For each motif a scheme illustration of the profile is presented. The X-axis represents taxonomy clades ordered by a descending evolutionary distance from human. The Y-axis represents the length normalized score for the protein versus each organism as compared to human.

### Global Motifs

Global motifs refer to the overall profile of a protein among all, or most, of the species. The global motifs are: *Steps, Critical, Lately Developed*, and *Plateau* (Fig. 3).

**Figure 3:**
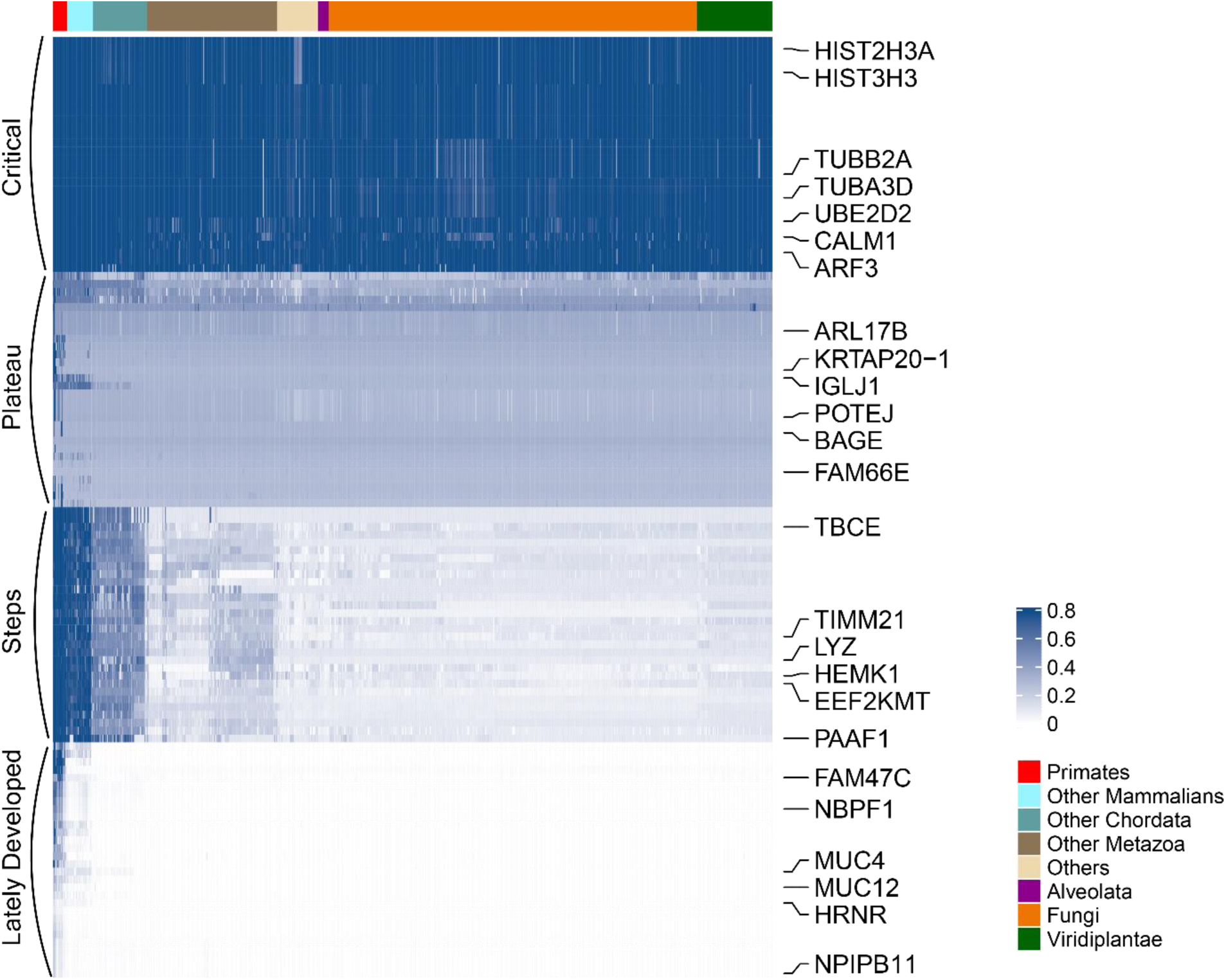
Gobal motif profiles heatmap. Heatmap of the length normalized profiles of four global conservation motifs. Rows are proteins and columns are taxonomy clades ordered by a descending evolutionary distance from human. Dark blue indicates that the protein is highly conserved within the species. The proteins are clustered with the Euclidean distance method and leaf reordering clustering method “ward”. The global *Steps, Critical, Lately Developed* and *Plateau* proteins are grouped into four distinct clusters.

#### Steps

The sequence similarity between two orthologs is a function of both the evolutionary distance between the species and the selective pressures that acted on those proteins. We would expect that, on average, proteins would behave as a “molecular clock” and would be more conserved in taxonomic clades closer to human, creating a *Step*-like pattern. This is indeed the case and our analyses generated a global *Steps* appearance where the protein scores in distant taxonomic clades were lower than those in closer taxonomic clades (i.e. the average sequence similarity score in Mammalians is higher than in Fungi, whose score is higher than in Viridiplantae). The proteins related to that motif form a stepwise behavior. The *Steps* motifs can also be local; primate proteins are more similar to human proteins as compared to orthologs from other chordata. Similarly, orthologs from the Mammalia clade are more conserved than orthologs from the rest of the animal kingdom.

Consequently, we wanted to identify which, and how many, proteins have global *Steps* behavior as a probable result of the molecular clock. To this end, we defined the sequence divergence of the human proteins in each organism and normalized it to the expected divergence. The expected divergence is the average difference between all the human proteins to their orthologs in each species. This means that the vector indeed gradually decreases when the species are ordered in a descending evolutionary distance from human, based on the NCBI Common Taxonomy Tree. We defined the *Steps* criterion to calculate the Euclidean distance of protein scores to the means vector (see Methods). This score indicates how close the protein is to the ‘average’. Overall, the *Steps* criterion distribution is very narrow, with an average distribution of 0.37 and a standard deviation of 0.23. In order to validate the significance of the results, we calculated the same *Steps* score on a shuffled version of the data (see Methods). The criterion distribution is very close to the shuffled distribution and to the “average protein” (Supplementary Fig. 1-A). This supports our aim of finding the most general motif among the protein sets that behave according to the expected evolutionary trend. Nevertheless, the criterion distribution is skewed to the right, which implies additional protein sets that respond to very different selective pressure and show different evolutionary patterns. We then turned to characterizing those protein sets that showed less trivial evolutionary patterns.

#### Critical

Some proteins are essential for sustaining life. These proteins show very low sequence variability among species and mutations within them and almost always show a reduction in organism fitness. If the function of these proteins is critical to the survival of the vast majority of species, we expect to find a *Critical* pattern. *Critical* proteins show a high sequence similarity, much more so than expected, among all eukaryotes (global) or in clades (local). These proteins are almost never lost in species.

We defined the *Critical* criterion to calculate the average sequence similarity in each of the 8 taxonomy clades and then summarized the means. We calculated the same score to a shuffled version of the data (see Methods). We defined proteins with scores higher than the maximal score in the shuffled version as C*ritical* (Supplementary Fig. 1-B). This revealed 548 *Critical* proteins (Supplementary Table 1.1). The average *Critical* protein exists (a highly conserved ortholog was found) in 48% of the species as compared to 6% of the species in the rest of the proteins. As expected, among the *Critical* proteins there was a significant enrichment of histones (hypergeometric test p-value: 4.88e-42), ribosomes (hypergeometric test p-value: 2.05e-67), actins (hypergeometric test p-value: 1.59e-10) and tubulins (hypergeometric test p-value: 8.83e-17). H3C15 (H3 Clustered Histone 15), H3-3A (H3.3 Histone A) and H3C1 (H3 Clustered Histone 1) are all ranked in the top of the *Critical* criterion.

It is well established that histone H3 proteins are highly conserved across all eukaryotes^22,23^. In comparison, H1 is known to be less conserved than other histones, probably because it is a linker histone^24^. We show that H1 is indeed ranked low in the *Critical* criterion. H2A and H2B, which are involved in regulation, are known to have sequence and structural variations and to be post-translationally modified^25^. Therefore, some of them are *Critical* while others are less conserved and ranked lower in the criterion. No variants are known for H4^26^ and they are highly ranked in the *Critical* criterion (Supplementary Fig. 2).

The next very *Critical* proteins are the actins, which are known to be highly conserved and are involved in various modes of cell motility and in the maintenance of the cytoskeleton^27^. Followed by the *Critical* protein groups of the α- and β- tubulins, which are also known to be highly conserved heterodimers, they act as structural components of microtubules, which are a cytoskeleton element of all eukaryotic cells^28^. An additional protein that emerged as *Critical* is the CALM1, Calmodulin Ca(2+)-sensing protein, which is known to be highly conserved in eukaryotes^29^.

The ‘nonsense-mediated mRNA decay’ GO annotation term is highly enriched within the *Critical* proteins (p-value: 1.17E-59) (Supplementary Table 2.1). ‘Nonsense-mediated mRNA decay’ is a crucial process regulating transcript quality and abundance and it is known to be conserved among eukaryotes^30–32^.

The ‘translational initiation’ GO annotation term is also highly enriched within the *Critical* proteins. ‘Translational initiation’ is a complex process that involves eukaryotic conserved initiation factors (such as EIF1B, EIF2S3, EIF4A2 and ribosomes) and regulation mechanisms that select the start codon on the mRNA^33^.

Interestingly, among the *Critical* proteins, some are not known to be highly conserved such as the ARL3 (ADP Ribosylation Factor Like GTPase 3) and the PRPF8 (Pre-MRNA Processing Factor 8), which are associated with Nonsyndromic Autosomal Dominant Retinitis Pigmentosa^34^.

#### Lately Developed

The evolution of species and taxonomic clades is frequently derived from new proteins that are unique to the clade. From the human perspective, most *Lately developed* proteins are absent among most eukaryotes and can be found specifically in primates and humans. These proteins are probably important for primate unique characteristics.

This protein set was identified in a complementary fashion to the *Critical* pattern (see Methods). Protein scores that are lower than the minimal score of the shuffled operation reflect the *Lately Developed* motif, which are proteins that are conserved less than expected randomly (Supplementary Fig. 1-C). Thirty-six proteins were identified as *Lately Developed* (Supplementary Table 1.1). Out of these were eight mucin proteins that were annotated as ‘innate immune response activating cell surface receptor signaling pathway’ (p-value: 8.26e-04) (Supplementary Table 2.2).

The mucins protein evolution is driven by a “Red Queen” effect, in which hosts are constantly in a race to evade the more rapidly evolving pathogens that infect them^35,36^. As the pathogens vary between different host species, the mucins proteins would vary as well.

Another enriched GO annotation is ‘establishment of skin barrier’ (p-value: 3.5e-05), which are proteins responsible for the epithelial barrier of the skin and for limiting its permeability. This term includes the FLG, HRNR and FLG2 proteins. Null mutations within the FLG gene encoding filaggrin have been identified in approximately 30% of atopic dermatitis patients, a common allergic skin disease^37^. HRNR differential expression levels have also been associated with atopic dermatitis susceptibility^38^. It is known that many aspects of the anatomy of the skin are unique to humans as compared to non-human mammals, and many drastic changes have occurred in the skin that allow thermoregulation^39^.

In addition, the NBPF protein family, which are part of the neuroblastoma breakpoint family, are ranked at the top of the *Lately Developed* criterion. Interestingly, this protein family has no discernible orthologues in rodent genomes and is the result of species-specific gene duplication events that occurred during primate evolution^40^. Some attempts have been made to connect the NBPF protein family to brain development^41^.

#### Plateau

*Plateau* proteins show a constant percentage of conservation at a specific value among most species. Such constant behavior is less expected as the conservation should, theoretically, change in correlation with the evolutionary distance from human. We defined the *Plateau* criterion to calculate the variance among the invertebrate clades and summarized the variances. In order to validate the significance of the results, we calculated the same score on permutations of the data (see Methods). For further analyses, we extracted the protein that scored lower than the minimal score in the shuffled operation as these proteins are “less random” than expected (Supplementary Fig. 1-D). This revealed 137 *Plateau* proteins (Supplementary Table 1.1).

We hypothesized that *Plateau* proteins contain highly conserved regions (in variable sizes) that dictate the percentage of conservation. Therefore we examined whether proteins that share the same constant conservation score tend to contain an equal sized structural domain. We determined the ‘Pfam coverage score’ as the length of the largest Pfam domain^42^ of a protein, divided by the protein length. We noticed that, in general, proteins Pfam-coverage-score-trend increased as the mean-conservation-score enlarged. In addition, the constant protein conservation value can be either unique or noisy. When comparing two *Plateau* proteins sets with different variance values, the Pfam coverage score among the low variance proteins was consistently larger than among the high variance proteins. This indicates that *Plateau* proteins with a higher variance include several smaller domains instead of a large conserved domain (Supplementary Fig. 3).

The *Plateau* motif provides information on the structure of a protein, based on its phylogenetic profile alone. As the information for the construction of the profile is based on 1096 species, the prediction is less biased by the choice of species for multiple sequence alignment. Furthermore, the *Plateau* motif answers the questions “what characterizes proteins that contain a large conserved domain” and “which highly conserved large domains are common”. For example, among the *Plateau* proteins there is a high representation of myosin proteins, which are a large family of motor proteins that share the common features of ATP hydrolysis, annotated by the ‘actin-myosin filament sliding’ GO annotation term (p-value: 3.72e-04, Supplementary Table 2.3). In addition, the POTE Ankyrin Domain family members are highly represented within the *Plateau* proteins under the ‘retina homeostasis’ GO annotation term (p-value: 0.02).

### Local motifs

The conservation and phylogenetic patterns of proteins can, and should, be described globally. Nevertheless, some “less trivial” evolutionary patterns have a more local nature, although fundamental to understanding protein function. Local motifs focus on proteins that show an unexpected profile in specific taxonomic groups. This includes loss or gain of the proteins at certain positions across the tree of life. Below, we describe the local motifs, which are: *Clade loss, Trait Loss*, and *Gain* (Fig. 2).

#### Loss

A loss event can occur at the common ancestor of a taxonomic clade causing a group of proteins to be absent from a monophyletic taxonomy clade. We termed this type of loss *Clade Loss*. Another possibility for a loss event is within species that share a common phenotype, which are part of a polyphyletic clade. We termed this type of loss *Trait Loss*. The hypothesis for both motifs is that the analyzed species share common characteristics that may provide an explanation for a significant loss of the proteins.

#### Clade Loss

*Clade Loss* proteins are significantly diverged or lost within a taxonomic clade, while conserved in close clades. *Clade Loss* proteins can describe many different and distinct evolutionary phenomena. One example is a protein group that is lost in invertebrate animals ‘Arthropoda’ but exists in ‘Viridiplantae’. We identified 102 proteins (Supplementary Table 1.1, see Methods) and found that 14 proteins were related to ‘steroid biosynthetic processes’ (p-value: 4.88e-10, Supplementary Table 2.4). It was previously reported that insects cannot synthesize cholesterol and other sterols and that they obtain sterols externally from plants^43^, which they then convert to cholesterol to ensure normal growth, development, and reproduction^44,45^. This insect *Clade Loss* motif helps explain observations with a simple rationale and analysis. The *Clade Loss* criterion can be easily applied to any taxonomic clade of interest, allowing the definition of any other lost individual or groups of proteins. The insect case study is an example of the ability of the *Clade Loss* motif to reveal informative biological insights from phylogenetic profiling data.

#### Trait Loss

One of the major challenges in biology is to associate genotype with phenotype. We and others have previously shown that protein phylogenetic profiles can be used to predict organism traits ^13,14,46,47^. *Trait Loss* proteins are unexpectedly diverged or lost within a polyphyletic clade, in correlation with certain traits. One example of the *Trait Loss* motif is metazoan parasites, which are a polyphyletic group of species. We used 69 metazoan human parasites in our analyses (Supplementary Table 1.2). We were interested in the proteins whose scores differed between these parasites and other metazoa (Fig. 4, see Methods).

**Figure 4:**
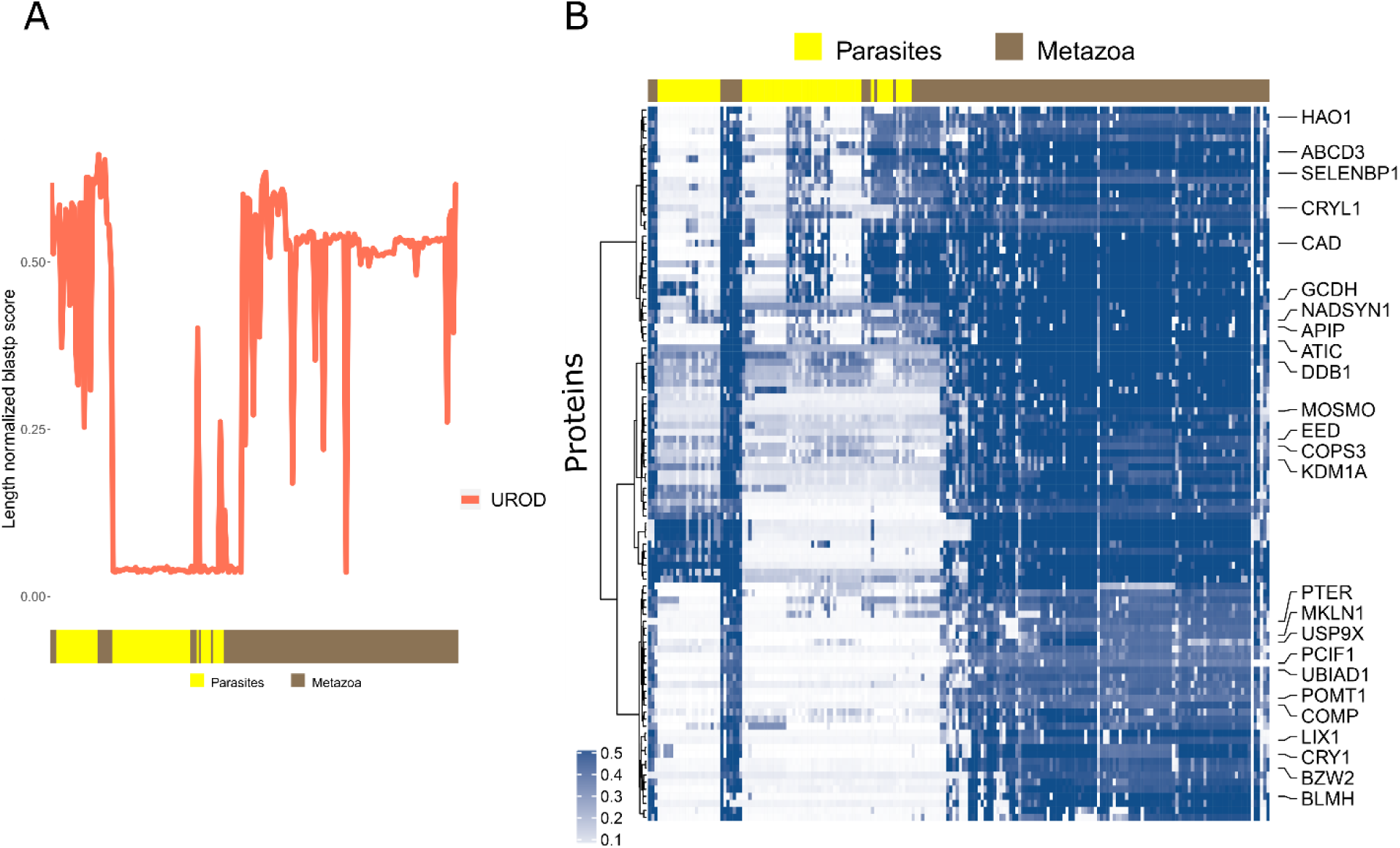
Local trait loss. (**A**) Example of a trait loss protein profile. Smoothed length normalized phylogenetic profiles of the clade loss protein UROD (Uroporphyrinogen Decarboxylase). The X-axis represents the metazoan taxonomy clade which includes the metazoan parasites (yellow) and other metazoa (brown). The Y-axis represents the length normalized blastp score fort the protein versus each organism as compared to human. A sudden absence of UROD is indicated specifically within the parasites and is considered as ‘trait lost’. (**B**) Heatmap of the trait lost proteins length normalized profiles among the metazoa taxonomic clade. Rows are 102 trait lost proteins. Columns are metazoan species including metazoan parasites (yellow) and other metazoa (brown). The proteins are clustered with the Euclidean distance method and leaf reordering clustering method “ward”. Dark blue indicates that the protein is highly conserved within the species. An absence of proteins is indicated specifically with the parasites.

Out of the 102 *Trait Loss* proteins (Supplementary Table 1.1), 35% were related to metabolism pathways that included amino acid metabolism (p-value: 9.05e-05) and glucose metabolism (p-value: 0.00612) (Supplementary Table 2.5). Several fatty acid catabolic proteins (e.g. HACL1, HAO1 and SCP2) disappeared from most metazoan parasites. Previous studies noted that tapeworms, like flukes, lacked the ability to synthesize fatty acids and cholesterol, and instead scavenged essential fats from their hosts^48–50^.

Interestingly, we found enrichment for peroxisome organization pathway proteins (p-value: 2.14e-06). The peroxisome is responsible for oxidative metabolism of lipids and the detoxification of reactive oxygen species. Previous work identified the loss of peroxisomes in other free living related species^51^. Members of the peroxisomal proteins are studied as a drug target for the development of novel therapies against diseases caused by parasites^52–55^. We propose that the peroxisomal protein set, absent specifically in metazoan parasites, can be considered as targets for drugs against human parasites.

In addition, the three proteins ALAD, UROD and CPOX, revealed in the motif and related to heme biosynthetic process, are also known to differ significantly between the human parasitic worms *B. malayi, O. volvulus*, and *W. bancrofti* and humans^56^. Previous analysis considered heme biosynthesis as a possible antimalarial target^57^. Other heme biosynthetic proteins such as HMBS, FECH and PPOX also observed a pattern of loss specific to metazoan parasites. These findings support the heme biosynthetic process proteins as valid targets for drugs against metazoan parasitic infections.

Finally, the COP9 signalosome (CSN) diverges in metazoan parasites where COPS6, GPS1, COPS3, DDB1, COPS4 are completely lost. The CSN regulates the cell cycle (with the involvement of ubiquitin machinery)^58,59^ and is involved in the regulation of apoptosis. Previous work has identified a CSN in *S. mansoni*^60^. We looked at differences in conservation in the caspases that interact with CSN and found reduced conservation in the effector/executor caspases 3,6 and 7^61^. This surprising result of a major diversion in the evolution of the CSN complex in metazoan parasites points to a possible alternative mode of action of this pathway in metazoan parasites. The importance of CSN in the regulation of proliferation and apoptosis, makes it a possible target for anti-parasite treatments.

#### Gain

Some patterns have local, highly conserved, orthologs that are found only in few distant organisms, while in other species there are no significantly similar orthologs. We termed this sporadic appearance of an ortholog, within a select few species, as *Gain*. A gain event at a monophyletic taxonomic clade can appear only in the reference proteome’s taxonomy clade, which in our case is the primate taxonomy.

The *Lately Developed* pattern is a specific case of that scenario. Therefore we focused only on gain in polyphyletic taxonomic clades. The rationale behind the *Gain* proteins is that their similarity within a few specific species does not correlate to the taxonomic distance of those species. Therefore we defined the *Gain* criterion to capture proteins whose conservation score, within a specific organism, is much higher than expected, considering the overall conservation scores of this organism’s proteome and the overall conservation of other species in its taxonomic clade.

We calculated the Z-scores of the Length Normalized Phylogenetic Profiling scores^13^, which measures how much a protein is conserved as compared to its expected conservation (see Methods). We defined the *Gain* criterion to detect gaps within the sorted Z-scores of each organism and focused on the top threshold proteins (see Methods). This criterion indicates an unexpected conservation score of a protein in a specific organism. We applied this criterion on the Fungi and Viridiplantae clade.

The *Gain* criterion revealed 40 proteins in the fungi taxonomy and 33 proteins in the Viridiplantae taxonomy (Supplementary Table 1.3). Among the *Gain* proteins in fungi is SEPSECS (O-phosphoseryl-tRNA(Sec) selenium transferase). Z-scores of this protein unexpectedly increased among the Smittium fungi clade in *S. culicis, S. simulii* and *S. angustum*, and among the Chytridiomycetes clade only in *G. prolifera*.

We validated the presence of a possible SEPSECS ortholog specific to these species by *blastp* the human SEPSECS versus all the Smittium species, and all the Chytridiomycetes species using the nr database. Significant results appeared only in the species where significant Z-scores appeared in our data (except for *S. megazygosporum* where *blastp* revealed a hypothetical protein of 30%) (Fig. 5-A, Supplementary Table 1.4).

**Figure 5:**
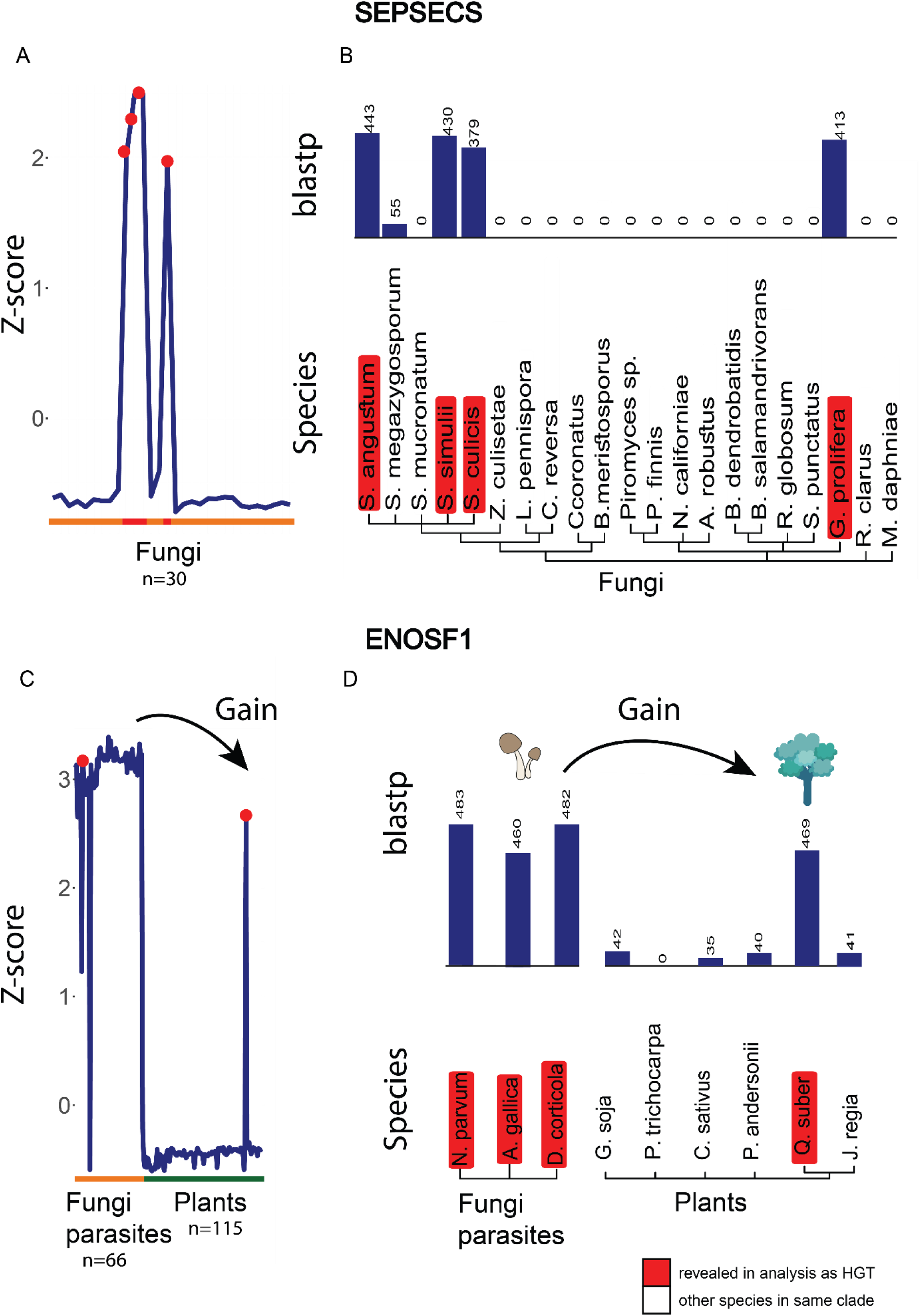
Examples of HGT events. (**A**) Smoothed Z-score phylogenetic profiles of the SEPSECS protein among the fungi taxonomic clade. The X-axis represents the fungi Zoopagomycota and Chytridiomycetes clade (orange). The Y-axis represents the Z-score of the SEPSECS in each organism. A sudden peak (red) of SEPSECS is observed at S. culicis, S. simulii, S. angustum and G. prolifera and is therefore a candidate for an HGT event. (**B**) Taxonomy tree of the Zoopagomycota and Chytridiomycetes taxonomic clades along with the blastp scores of the SEPSECS protein in each organism. Annotaded in red are the species where SEPSECS was revealed as gained within them according to the gain criterion. (**C**) Smoothed Z-score phylogenetic profiles of the ENOSF1 protein among the fungi and plants taxonomic clade. The X-axis represents the Dothideomycetes and Agaricales fungi (orange) and Fabids plants (green) taxonomic clades. The Y-axis represents the Z-score of ENOSF1 in each organism. A sudden peak (red) of ENOSF1 is observed at Q. suber and its parasites and is therefore a candidate for an HGT event. (**D**) Taxonomy tree of the Fabids clades and Q. suber parasites along with the blastp scores of the ENOSF1 protein in each organism. The Q. suber and its parasites are annotated in red because ENOSF1 was revealed as gained within them according to the gain criterion.

Another protein that was revealed in the *Gain* criterion in plants is ENOSF1 (Mitochondrial enolase superfamily member 1), which showed a sudden peak in *Q. suber*, a cork oak part of the *fabids* taxonomy. We validated the presence of a possible ENOSF1 ortholog specific to *Q. suber* by *blastp* the human ENOSF1 versus candidate species in each of the fabids sub-clades. No significant score was obtained in any of the species except for *Q. suber* (Fig. 5-B, Supplementary Table 1.4).

A sudden appearance of such proteins in specific species may suggest horizontal gene transfer (HGT) events. SEPSECS is involved in a two-step pathway with PSTK (Phosphoseryl-TRNA Kinase), which is mentioned in the literature as laterally transferred to fungi^62^. The NPP profile of ENOSF1 among all eukaryotes is extremely variable. An unexpected phylogenetic distribution of this protein in bacterial lineages has been reported^63^. Most of the bacteria and fungi that have the ENOSF1 protein are human, animal and/or plant pathogens. Three of the fungi pathogens known to affect *Q. suber* that exist in our data set are *D. corticola, A. gallica* and *N. parvum*^64^. We confirmed that they all have the ENOSF1 protein by their NPP Z-scores, and by *blastp* the ENOSF1 human protein versus those genomes. This observation strongly suggests a horizontal gene transfer event from these pathogens to the *Q. suber*.

The *Gain* motif revealed possible proteins and species candidates for horizontal gene transfer events. The *Gain* criterion on the fungi and plants taxonomy revealed 73 overall possible candidates for HGT events along with the candidate species (Supplementary Table 1.3).

## Discussion

Protein evolution is an extremely complex and variable process that is affected by different evolutionary forces, such as positive and purified selection, drift, gene flow and even HGT. These events vary in every species, affecting an organism’s fitness and driving its evolution. Despite this complexity, the scientific community currently uses a relatively limited vocabulary, lacking in dimensionality, when describing protein evolution along different species. This definition of conserved, poorly conserved, etc., seems rather simplistic when aiming to describe and study a subject as complex as protein evolution across many species. By identifying and describing common conservation patterns, “conservation motifs”, we were able to enrich the language for describing and studying protein evolution. Additionally, we were also able to identify proteins with unexpected patterns of conservation. This flexible terminology makes it possible to characterize and, for the first time, communicate protein conservation patterns simply and more accurately.

Here, we analyzed the evolutionary profiles of 20,294 human proteins against 1096 species and offered a novel terminology of the observed protein evolution. We identified seven conservation motifs: *Steps, Critical, Lately Developed, Plateau, Clade Loss, Trait Loss* and *Gain*, which present both expected and more complex evolutionary patterns. We provide a richer but simpler language than the basic terminology of “conserved or not conserved”. This study highlighted and defined common conservation patterns, associating them with protein evolution and function. Overall conservation motifs revealed biological and evolutionary insights about protein function and interaction with the environment.

Furthermore, we believe that this terminology is applicable for describing the evolution of most human proteins better than the terms currently in use. For example, TP53 (Tumor Protein P53), the most frequently mutated gene in human cancer, has a *Steps* motif and is not conserved in all the non-metazoan eukaryotes (Fig. 6-A). ACTA2 (Actin Alpha 2), a highly conserved protein, observes a *Critical* motif (Fig. 6-B). CDK4 (Cyclin Dependent Kinase 4) is *Steps* until Metazoa and *Plateau* in the rest of the eukaryotes. A *Local Loss* of conservation is indicated within the Metazoa clade (Fig. 6-C).

**Figure 6:**
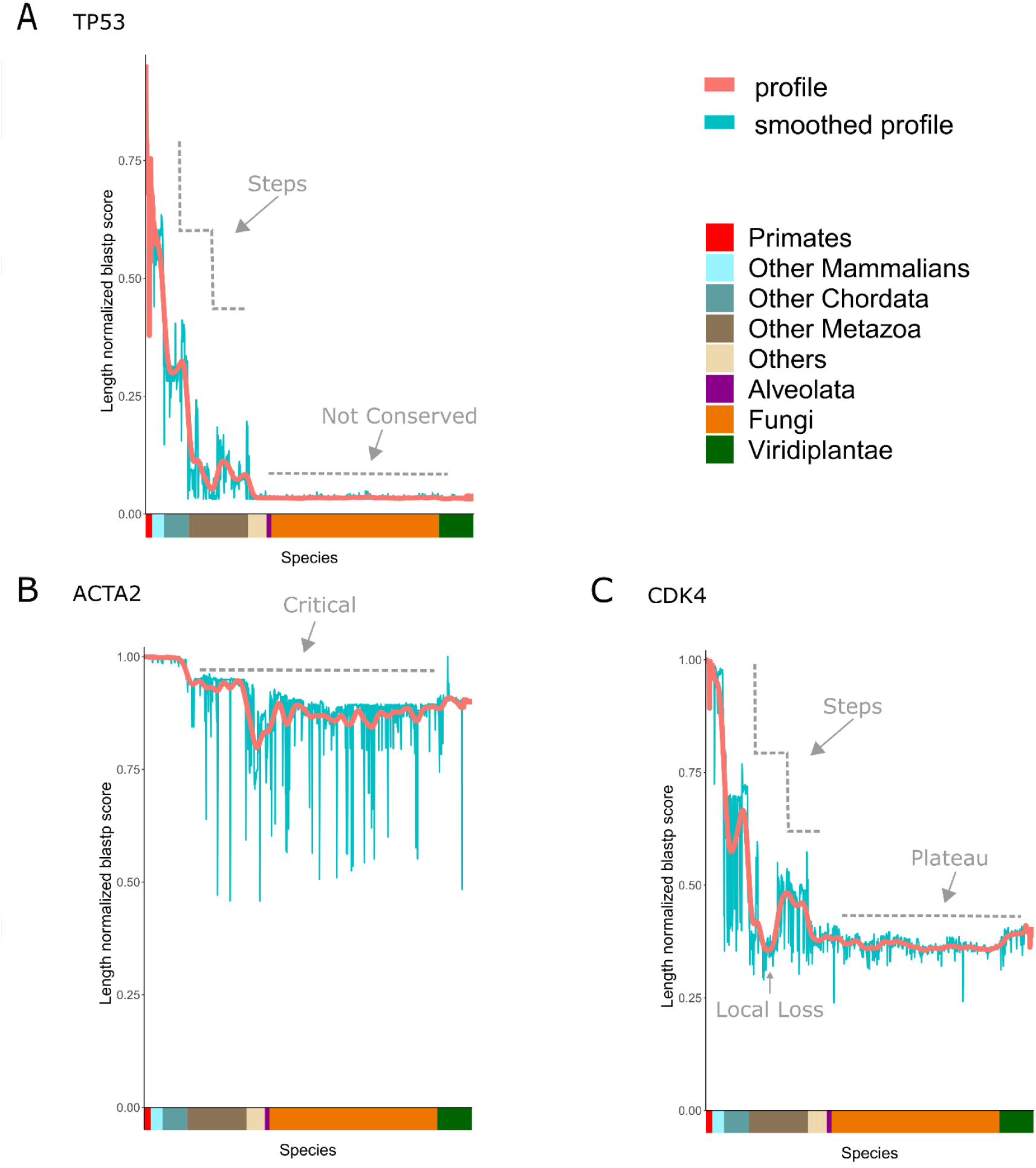
Example of highly-sited protein conservation motifs. Length normalized phylogenetic profiles along with a smoothed line of three proteins and their conservation motif definition. (**A**) profile of *Steps* TP53. (**B**) profile of *Critical* ACTA2. (**C**) profile of CDK4 which is local *Steps* up until metazoa and local *Plateau* in the rest of eukaryotes. The X-axis represents the 8 taxonomic clades ordered by a descending evolutionary distance from human. The Y-axis represents the length normalized blastp score of the proteins versus each organism as compared to human. Light-blue line represents the actual scores and the pink line represents the smoothed scores.

Conservation motifs can shed light on evolutionary processes of proteins of interest. For example, it can point to the high divergence of the mucins - a family of heavily glycosylated proteins that are produced in most animals to form gels. These evolved rapidly and present a *Lately Developed* motif. Obviously, conservation motifs can predict insights about proteins, such as structure and domain coverage, revealed from the *Plateau* motif. Moreover, conservation motifs can relate phenotype to genotype such as to a protein group absent exclusively among polyphyletic parasites (*Trait Loss*). The discovery of this behavior provides a clue as to the function of these proteins as well as to highlight them as candidate targets for drug development.

Conservation motifs also revealed a set of 73 candidate proteins for horizontal gene transfer events, along with a hint about the organisms where the transfer occurred. Furthermore, conservation motifs explain evolutionary relations between close and distant species. One example is the absence of sterol proteins among insects, but which is compensated by their presence in plants and points to strong interactions between these two taxonomic groups. Another example is the ability to note the donor and acceptor organisms of horizontal gene transfer events and to predict the putative proteins. Specifically, we presented a lateral transfer of ENOSF1 to *Q. suber* from its fungal parasites. Of 18 proteins revealed as horizontally transferred to *Z. mays*, seven of these proteins are paralogues. Further investigation is needed to better understand the lateral transfer process to *Z. mays*.

One current limitation of this analysis results from poor genome assembly that are incomplete. This causes false negatives since the absence of proteins is not fully proven. Therefore, when detecting protein absence, the biological hypotheses revealed from the patterns should be carefully considered in order to avoid mistaken insights. For example, the reason for the significant loss of proteins among the ‘Aves’ class is probably a result of a high GC content in bird genomes and not an actual signal of protein absence^65^.

As an annotation or language, we believe that this work will open a new and more relevant way to consider conservation patterns. Following the exponential growth in the number of sequenced genomes, the annotation can be improved and expanded to include other motifs. We hope that over time, the biological meaning of these motifs will be better understood. For example, we chose the *Arthropoda* and *Viridiplantae* taxonomic clades to demonstrate the *Clade Loss* motif, and the parasites species as an example of polyphyletic species (where the proteins are absent) to demonstrate the *Trait Loss* motif. However, this analysis can and should be applied to any taxonomic group of interest.

In conclusion, there is an unmet need to describe protein evolution in simple terms. As far as we are aware, this work is the first to define a method for describing protein evolution terminology. During an era of exponential growth in the genomic data, we believe that this work offers a new language for proteins and can enhance our understanding of their evolution.

## Methods

### Creating the Length Normalized Phylogenetic Profiling Matrix

#### Data retrieval

The protein phylogenetic profiling data was generated as previously described^13,14,21^. 1154 proteomes were downloaded from both NCBI’s RefSeq non-redundant protein database and Uniprot. The script ‘update_blastdb.pl’ was used for the download of the non-redundant protein database^66^ (accessed at 25.12.2018, link - ftp://ftp.ncbi.nlm.nih.gov/blast/db/). In order to extract 1154 specific taxonomies from the compressed RefSeq downloaded database, we used the ‘taxonkit’ tool, which maps the sequence identifiers to the relevant species and extract the relevant fasta files. The proteomes from Uniprot were downloaded as FASTA files using the REST API (The UniProt Consortium 2019) (accessed at 28.12.2018, link - https://www.uniprot.org/uniprot/?query=proteome:Proteome_ID&format=fasta).

The reference human proteome was retrieved from UniProt as well (accessed at 28.12.18) and filtered such that each gene had a single representative protein when the largest protein for each gene was considered. We reached 20294 human reference proteins.

#### Length normalized score calculation

For each of the human proteins (20294) we calculated a continuous score (between 0 to 1) that reflects its quantitative conservation among the large set (1154) of eukaryotic proteomes. The continuous score was calculated by:

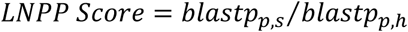

*blastp*_*p,s*_ = *blastp* bit score of the human query protein *p* aligned with its one-directional best hit in one of the 1154 subject proteomes *s*.

*blastp*_*p,h*_ = *blastp* bit score of the human query protein *p* aligned with its one-directional best hit in the human proteome *h*.

Since the bit score depends on the length of the protein, this score reveals a length normalized score of the protein conservation.

We executed *blastp* on the command-line^67^ (version 2.7.1) with the argument ‘-max_target_seqs 1’ to retain only the top hit per protein per species.

The resulting matrix *M* is 20294 rows (proteins) x 1154 columns (species), where *M*_*i,j*_= blastp score of protein *i* in organism j, divided by protein length. The matrix is called LNPP-Length Normalized Phylogenetic Profiling. Most of the conservation motifs criteria are based on the LNPP matrix.

#### Z-Score calculation

For the *Gain* motif we used the Z-Score Normalized Phylogenetic profiling data^13,14^. For each of the human proteins (20294) we calculated the Z-Score that reflects how much the protein is conserved in (1154) eukaryotic proteomes more than expected. The Z-Score was calculated based on the LNPP matrix:

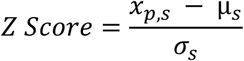

*x*_*p,o*_ = length normalized phylogenetic profiling score of protein *p* in proteome *s*

*μ*_*s*_ = the mean of all the proteins in proteome *s*

*σ*_*s*_ = the standard deviation of all the proteins in this proteome *s*.

#### Species selection

Some of the LNPP columns, representing 1152 species, were excluded from the analysis in order to reduce noise. First, we selected only one representative organism for each taxonomy strain, avoiding duplicated columns that might bias the analysis. Second, we detected some species whose scores among all the proteins is very low. This is a result of small proteome size or bad proteome annotation. Therefore we calculated the mean of each organism among all the proteins and excluded from the analysis the bottom 5% species. We considered a final set of 1096 species for the motifs identification (Supplementary Table 1.5).

#### Taxonomy Clades reordering

The species taxonomy kingdoms were ordered in a descending evolutionary distance from homo sapiens. Furthermore, we reordered the internal species within the taxonomy clades, in order to place closer species next to each other. The information about the species order was downloaded from the NCBI Common Taxonomy Tree architecture (link-https://www.ncbi.nlm.nih.gov/Taxonomy/CommonTree/wwwcmt.cgi) of the 1096 species. The tree was then rerooted for homo sapiens using the command nw_reroot in the Unix shell, belong to the newick utilities 1.6 tool (link-http://cegg.unige.ch/newick_utils). The tree was then reordered using the R generic ‘reorder’ function (belongs to the ‘stats’ package) using the ‘OLO’ method, based on the Pearson correlation of the species LNPP scores.

### Conservation motifs definition

#### Mathematical criteria

For all the mathematical criteria that included the taxonomy clades in calculations, we considered 8 taxonomy clades: Primates, Other Mammalians, Other Chordata, Other Metazoa, Alveolata, Fungi, Viridiplantae and Others, so that *n* = 8 consistently.

#### Steps criterion

*Euclidean distance to the means vector*.

For each protein *p* we calculated the score:

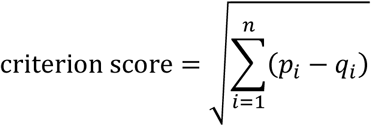

*p =* (*p*_1_, *p*_2_,…, *p*_*n*_)

*q =* (*q*_1_, *q*_2_,…, *q*_*n*_)

*p*_*i*_ *=* mean of length normalized score of protein *p* in clade *i*.

*q*_*i*_ = mean of length normalized score of all the proteins in clade *i*.

We ordered the proteins in a *descending* order of the criterion scores. In addition, in order to validate the significance of the results, we applied the above criterion on a shuffled version of the data, i.e. shuffling the protein means among each clade *i*.

#### Critical criterion

*sum of clades means*.

For each protein *p* we calculated the score:

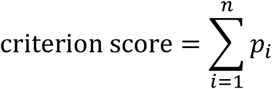

*p*_*i*_ *=* mean of length normalized score of protein *p* in clade *i*.

Since the 8 taxonomy clades sizes are variable, we summarized the means and all of the scores so that all clades would have the same power to affect the score. We ordered the proteins in a *descending* order of the criterion scores. In addition, in order to validate the significance of the results, we applied the above criterion on a shuffled version of the data by shuffling the protein means among each clade *i*. We extracted proteins that received a score higher than the maximal value in the shuffling operation after the shuffling was simulated 1000 times, and then the maximal value of all iterations was picked (maximal value: 5).

#### Lately Developed criterion

*sum of clades means*.

For each protein *p* we calculated the score:

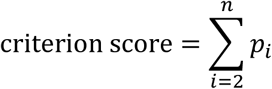

*p*_*i*_ *=* mean of length normalized score of protein *p* in clade *i*.

We ordered the proteins in an *ascending* order of the criterion scores. We did not include the Primate taxonomy clade in this criterion, since we were interested in proteins that were absent among most eukaryotes except for primates. Therefore we want the score to be minimal in all eukaryotes except for primates (7 clades). In addition, in order to validate the significance of the results, we applied the above criterion on a shuffled version of the data, i.e. by shuffling the protein means among each clade *i*. We extracted proteins that received a score lower than the minimal value in the shuffling operation after the shuffling was simulated 1000 times, and then the minimal value of all iterations was picked (minimal value: 0.21).

#### Plateau criterion

*distance of means from Plateau values*.

For each protein *p*, with a mean of length normalized score greater than 0.3, we calculated the score:

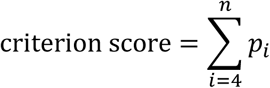

*p*_*i*_ *=* variance of length normalized score of protein *p* in clade *i*.

Some of the proteins that had a constant percentage of conservation were a result of very low conservation and short sequence. In order to avoid their emergence, we limited the protein group in advance to proteins that had an average length normalized score greater than 0.3. We did not include in the criterion the Vertebrates taxonomic clades, since within them most of the proteins show pattern of S*teps*. In addition, in order to validate the significance of the results, we applied the above criterion to a shuffled version of the data, i.e. by shuffling the protein variance among each clade *i*. We extracted proteins that received a score lower than the minimal value in the shuffling operation, when the shuffling was simulated 1000 times, and then the minimal value of all iterations was picked (minimal value: 0.008). The protein group that emerged included some *Critical* proteins, which we did not include in the final set of the *Plateau* proteins.

### Loss

The threshold used for these motifs to cut the proteins ranked in the criterion was 0.5%, thus limiting the motif size to ∼100 ranked proteins.

#### Clade Loss criterion

*distance of scores in two monophyletic clades*.

For example, for Arthropoda and Viridiplantae clades, we calculated the score for each protein *p*:

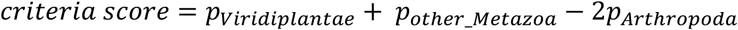

*p*_*Viridiplantae*_ = mean score of protein *p* in Viridiplantae clade

*p*_*Arthropoda*_ = mean score of protein *p* in Arthropoda clade

*p*_*other_Metazoa*_ = mean score of protein *p* in the rest of the species in the Metazoa clade

We ordered the proteins in a *decreasing* order of the criterion scores and picked the top threshold.

#### Trait Loss criterion

*distance of scores in two polyphyletic clades*.

For example for Metazoa and parasitic Metazoa, we calculated the score for each protein *p*:

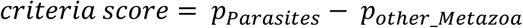

*p*_*Parasites*_ = mean score of protein *p* in a set of metazoan parasitic species.

*p*_*other_Metazoa*_ = mean score of protein *p* in the rest of the species in the Metazoa clade.

We ordered the proteins in a *decreasing* order of the criterion scores and picked the top threshold.

### Gain

#### Gain criterion

*distance between two following z-scores*.

For example: clade fungi. For each protein *p* we calculated the score:

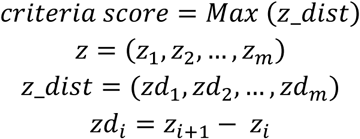

*z* = the z-score of protein p in each of the fungi clade species.

*z_dist* = the distance between two following z-scores. m = number of species in fungi clade

We ordered the proteins in a *decreasing* order of the criterion scores and picked the top threshold. To reduce noise, we further limited the criterion for proteins where their maximal gap between the sorted z-scores appear at the top quartile of the values, i.e. the unexpected high conservation score exists in no more than 25% of the clade species.

### Pfam domains retrieval

We downloaded the data about Pfam domains using the ‘biomaRt’ R package, using the ‘hsapiens_gene_ensembl’ dataset and ‘ensembl’ mart. Using the getBM function, we queried for the attributes: ‘external_gene_name’, ‘pfam’, ‘pfam_start’, ‘pfam_end’, for the filters: ‘hgnc_symbol’ where the values are all the NPP protein names.

### Phylogenetic profiling smoothing visualization

In order to illustrate specific phylogenetic profiles of proteins, we generated a geometric line where the x-axis represents all eukaryotes available in the time of the analysis, annotated to 8 taxonomy clades, and the y-axis is the normalized score that the protein received versus each specie compared to human. In order to reduce noise in visualization, smoothing methods were applied on the geometric line using the ‘smth’ function in the ‘smoother’ R package, with a window of 0.05 and a ‘Gaussian’ method.

### Statistical analyses

In order to calculate the significance of the presence of Histone, Ribosomes and Tubulins within the *Critical*s proteins, we used a hypergeometric test, using the R function ‘phyper’ belonging to the ‘stats’ package. For example, in order to calculate the significance of the presence of the histones proteins within the criterion proteins, the p-value is the probability of getting k or more histones in a sample of size, n, where the population contains m histones and RNPP-m other proteins.

K= the length of intersection of histones proteins and NPP proteins.

N= number of *Critical* proteins, number of trials.

M= number of histones proteins in NPP.

RNPP= all the proteins in the NPP.

### Enrichment analyses

We applied enrichment analysis on the conservation motifs protein in order to further assess the terms highly enriched within the protein groups. The enrichment was done using the function ‘gprofiler’ belonging to the ‘gProfileR’ R package. The parameters of the function are: organism = “hsapiens”, src_filter=“GO:BP”, correction_method=“fdr”, max_set_size= 200, min_set_size=15, min_isect_size=3, and max_p_value=0.05. We observed that many of the terms contain the same proteins and that they basically represent the same biological process. Therefore, we filtered the terms by extracting those that had more than 90% overlapping proteins, by considering terms with better p-values.

## Supplementary Figures

**Figure S1:**
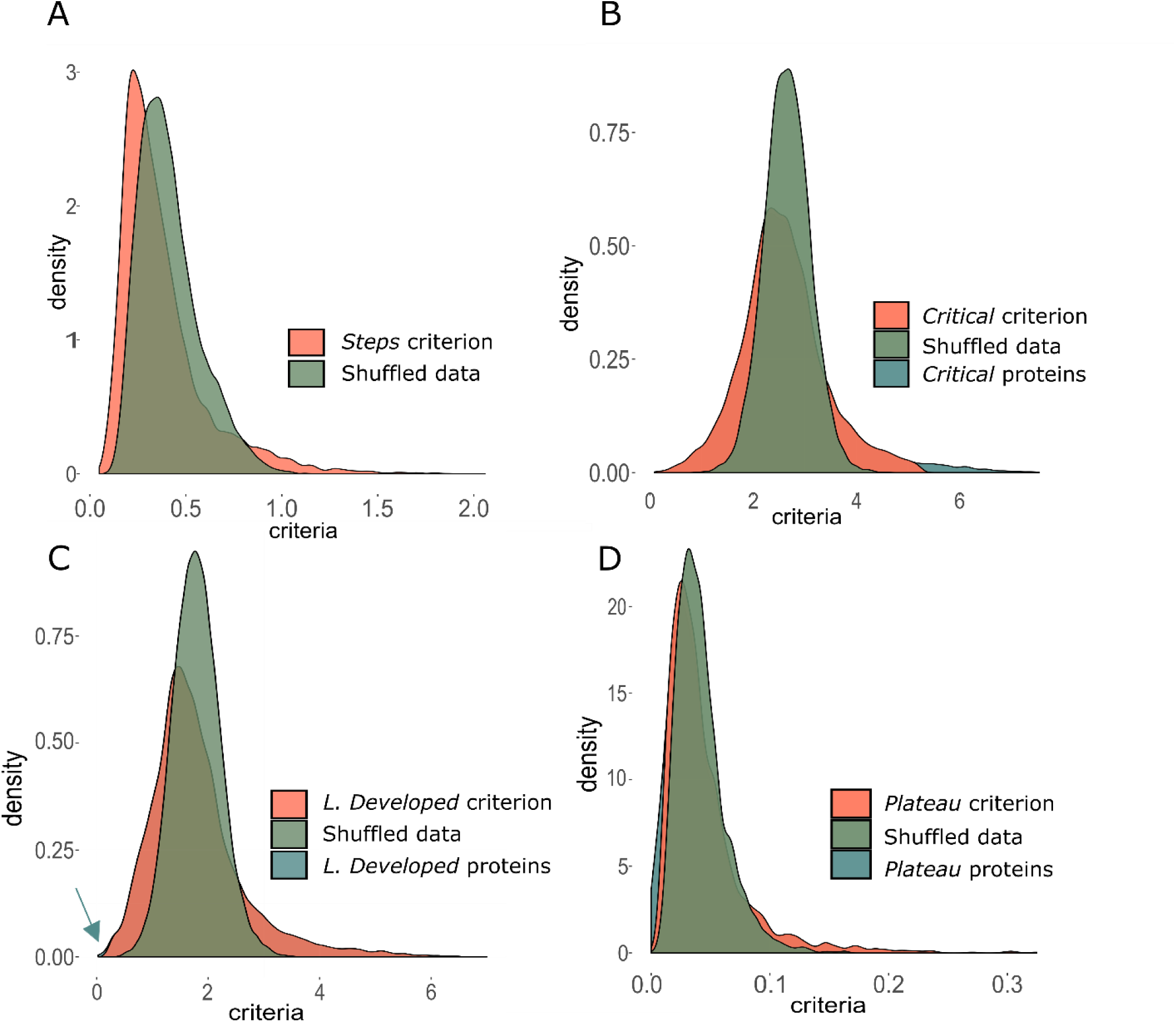
Distribution plots of the global conservation motif criteria. The X-axis represents the protein criteria scores. The Y-axis represents the density. The motif criteria (pink) are illustrated versus the same score when applied to shuffled data (green). The proteins with a score higher/lower than the maximal/minimal value of the criteria when applied to shuffled data were considered as ‘conservation motif’ proteins (blue). (**A**) Distribution plots of the *Steps* motif criterion. (**B**) Distribution plots of *Critical* motif criterion. (**C**) Distribution plots of the *Lately Developed* motif criterion. (**D**) Distribution plots of the *Plateau* motif criterion.

**Figure S2:**
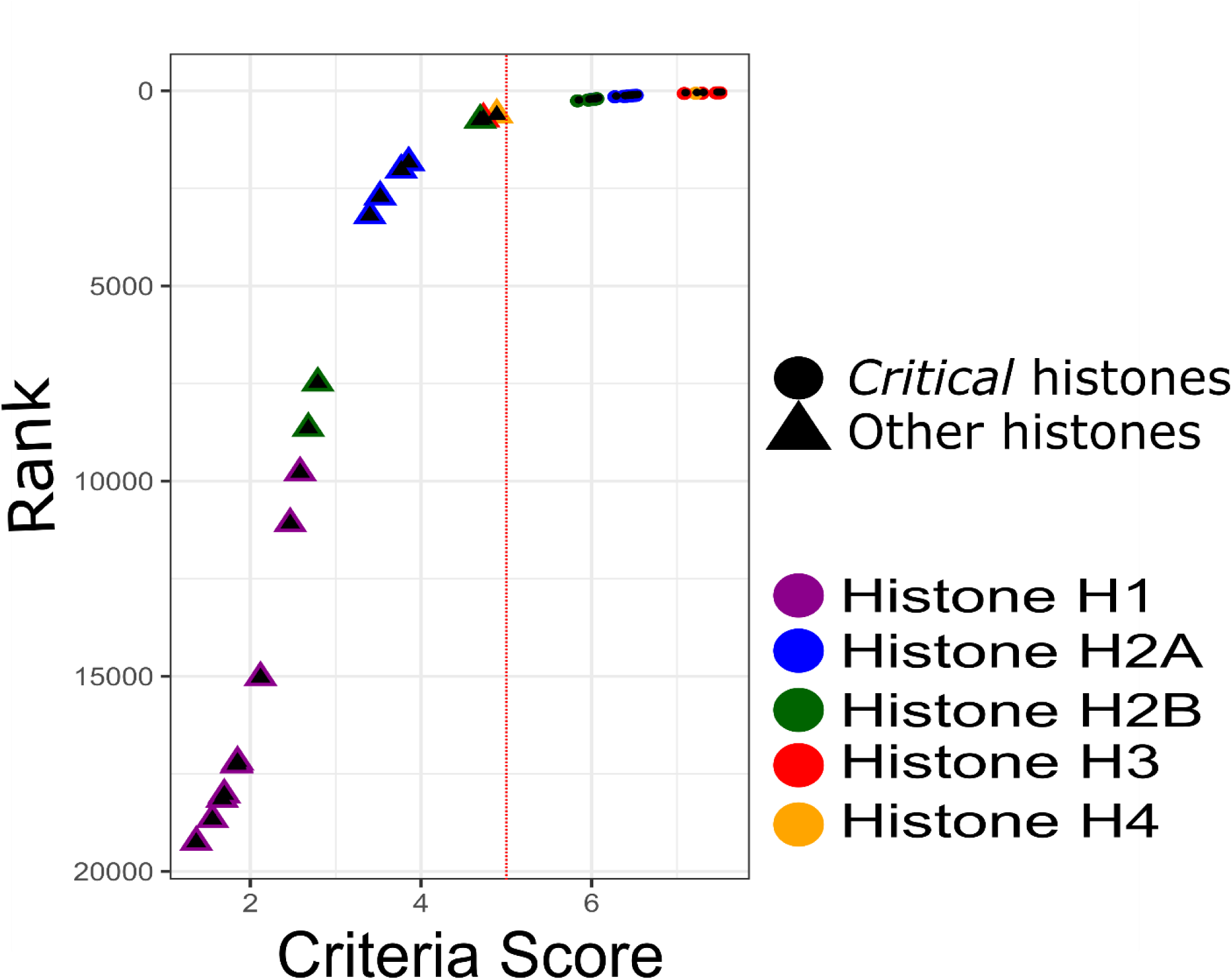
Histones rank in *Critical* criterion. Illustration of the different histones scores versus their rank in the critical criterion. The X-axis represents the histones scores in the *Critical* criterion, and the Y-axis represents the rank of the histones in the criterion. The different types of histones: H1, H2A, H2B, H3, and H4 are colored separately. Circles are histones that were defined as *Critical*s according to the *Critical* criterion. Triangles are histones that were not defined as *Critical*s according to the *Critical* criterion.

**Figure S3:**
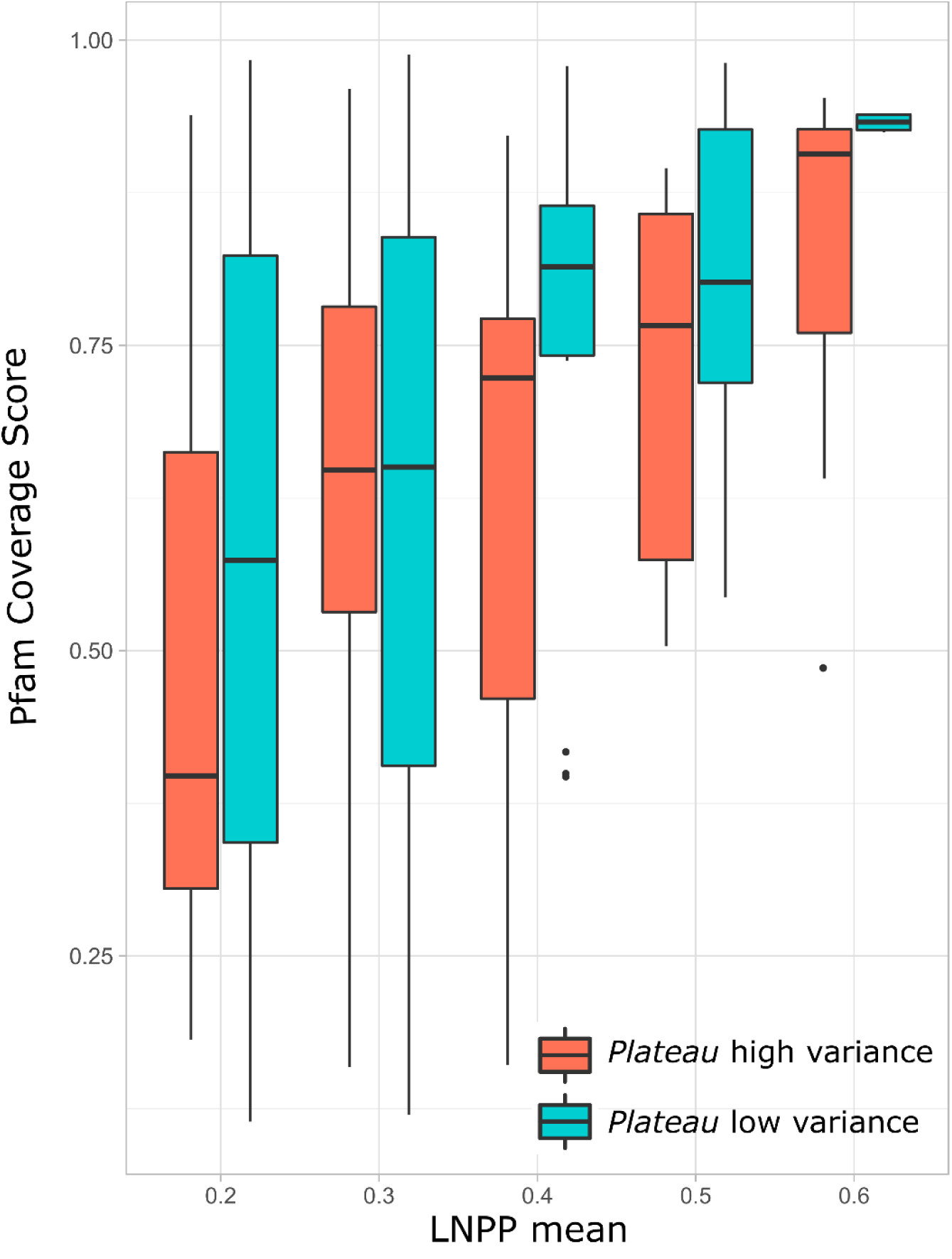
Pfam coverage distribution. Distribution of Pfam coverage scores for different means of length normalized scores. The X-axis represents means of length normalized scores, binned into 0.1 ranged intervals. The Y-axis represents the Pfam coverage scores. The red box plots are *Plateau* proteins with high variance while the blue box plots are *Plateau* proteins with low variance.

## References

1. Alon, U. Network motifs: Theory and experimental approaches. Nature Reviews Genetics 8, 450–461 (2007).

2. Han, J. D. J. et al. Evidence for dynamically organized modularity in the yeast protein-protein interaction network. Nature 430, 88–93 (2004).

3. Ashburner, M. et al. Gene ontology: Tool for the unification of biology. Nature Genetics 25, 25–29 (2000).

4. Bairoch, A. The ENZYME database in 2000. Nucleic Acids Res. 28, 304–305 (2000).

5. Radivojac, P. et al. A large-scale evaluation of computational protein function prediction. 10, (2013).

6. Thul, P. J. et al. A subcellular map of the human proteome - Supplemental material. Science (80-.). 356, eaal3321 (2017).

7. Uhlén, M. et al. Tissue-based map of the human proteome. Science (80-.). 347, (2015).

8. Papatheodorou, I. et al. Expression Atlas: gene and protein expression across multiple studies and organisms. Nucleic Acids Res. 46, D246–D251 (2018).

9. Aebersold, R. et al. How many human proteoforms are there? Nature Chemical Biology 14, 206–214 (2018).

10. Mitchell, A. L. et al. InterPro in 2019: improving coverage, classification and access to protein sequence annotations. Nucleic Acids Res. 47, D351–D360 (2019).

11. Cheng, H. et al. ECOD: An Evolutionary Classification of Protein Domains. PLoS Comput. Biol. 10, e1003926 (2014).

12. Pellegrini, M., Marcotte, E. M., Thompson, M. J., Eisenberg, D. & Yeates, T. O. Assigning protein functions by comparative genome analysis: Protein phylogenetic profiles. Proc. Natl. Acad. Sci. U. S. A. 96, 4285–4288 (1999).

13. Tabach, Y. et al. Identification of small RNA pathway genes using patterns of phylogenetic conservation and divergence. Nature 493, 694–8 (2013).

14. Tabach, Y. et al. Human disease locus discovery and mapping to molecular pathways through phylogenetic profiling. Mol. Syst. Biol. 9, 692 (2013).

15. Findlay, S. et al. SHLD 2/FAM 35A co-operates with REV 7 to coordinate DNA double-strand break repair pathway choice. EMBO J. 37, (2018).

16. Malcov-Brog, H. et al. UV-Protection Timer Controls Linkage between Stress and Pigmentation Skin Protection Systems. Mol. Cell 72, 444-456.e7 (2018).

17. Nordlinger, A. et al. Mutated MITF-E87R in Melanoma Enhances Tumor Progression via S100A4. J. Invest. Dermatol. 138, 2216–2223 (2018).

18. Omar, I. et al. Schlafen2 mutation in mice causes an osteopetrotic phenotype due to a decrease in the number of osteoclast progenitors. Sci. Rep. 8, (2018).

19. Sadreyev, I. R., Ji, F., Cohen, E., Ruvkun, G. & Tabach, Y. PhyloGene server for identification and visualization of co-evolving proteins using normalized phylogenetic profiles. Nucleic Acids Res. 43, W154–W159 (2015).

20. Schwartz, S. et al. High-Resolution mapping reveals a conserved, widespread, dynamic mRNA methylation program in yeast meiosis. Cell 155, 1409–1421 (2013).

21. Sherill-Rofe, D. et al. Mapping global and local coevolution across 600 species to identify novel homologous recombination repair genes. Genome Res. 29, 439–448 (2019).

22. Waterborg, J. H. Evolution of histone H3: emergence of variants and conservation of post-translational modification sites. Biochem. Cell Biol. 90, 79–95 (2012).

23. Talbert, P. B., Meers, M. P. & Henikoff, S. Old cogs, new tricks: the evolution of gene expression in a chromatin context. Nature Reviews Genetics 20, 283–297 (2019).

24. Henikoff, S. & Smith, M. M. Histone variants and epigenetics. Cold Spring Harb. Perspect. Biol. 7, (2015).

25. Shaytan, A. K., Landsman, D. & Panchenko, A. R. Nucleosome adaptability conferred by sequence and structural variations in histone H2A-H2B dimers. Current Opinion in Structural Biology 32, 48–57 (2015).

26. Pusarla, R. & Bhargava, P. Histones in functional diversification. FEBS J. 272, 5149–5168 (2005).

27. Pollard, T. D. What we know and do not know about actin. Handb. Exp. Pharmacol. 235, 331–347 (2017).

28. Gadadhar, S., Bodakuntla, S., Natarajan, K. & Janke, C. The tubulin code at a glance. J. Cell Sci. 130, 1347–1353 (2017).

29. Halling, D. B., Liebeskind, B. J., Hall, A. W. & Aldrich, R. W. Conserved properties of individual Ca2+-binding sites in calmodulin. Proc. Natl. Acad. Sci. U. S. A. 113, E1216–25 (2016).

30. Causier, B. et al. Conservation of Nonsense-Mediated mRNA Decay Complex Components Throughout Eukaryotic Evolution. Sci. Rep. 7, (2017).

31. Tian, M. et al. Nonsense-mediated mRNA decay in Tetrahymena is EJC independent and requires a protozoa-specific nuclease. Nucleic Acids Res. 45, 6848–6863 (2017).

32. Longman, D. et al. DHX34 and NBAS form part of an autoregulatory NMD circuit that regulates endogenous RNA targets in human cells, zebrafish and Caenorhabditis elegans. Nucleic Acids Res. 41, 8319–31 (2013).

33. Schmitt, E., Coureux, P. D., Monestier, A., Dubiez, E. & Mechulam, Y. Start codon recognition in eukaryotic and archaeal translation initiation: A common structural core. International Journal of Molecular Sciences 20, (2019).

34. Fahim, A. T., Daiger, S. P. & Weleber, R. G. Nonsyndromic Retinitis Pigmentosa Overview Clinical Characteristics of Nonsyndromic Retinitis Pigmentosa Clinical Manifestations of Retinitis Pigmentosa. GeneReviews (2020).

35. Varki, A. Evolutionary forces shaping the Golgi glycosylation machinery: why cell surface glycans are universal to living cells. Cold Spring Harb. Perspect. Biol. 3, (2011).

36. Cross, B. W. & Ruhl, S. Glycan recognition at the saliva – oral microbiome interface. Cellular Immunology 333, 19–33 (2018).

37. Park, K. D., Pak, S. C. & Park, K. K. The pathogenetic effect of natural and bacterial toxins on atopic dermatitis. Toxins 9, (2017).

38. Eaaswarkhanth, M. et al. Atopic dermatitis susceptibilityvariants infilaggrin hitchhike hornerin selective sweep. Genome Biol. Evol. 8, 3240–3255 (2016).

39. Rittié, L. Cellular mechanisms of skin repair in humans and other mammals. Journal of Cell Communication and Signaling 10, 103–120 (2016).

40. Vandepoele, K., Van Roy, N., Staes, K., Speleman, F. & van Roy, F. A novel gene family NBPF: intricate structure generated by gene duplications during primate evolution. Mol. Biol. Evol. 22, 2265–74 (2005).

41. Fiddes, I. T., Pollen, A. A., Davis, J. M. & Sikela, J. M. Paired involvement of human-specific Olduvai domains and NOTCH2NL genes in human brain evolution. Hum. Genet. 138, 715–721 (2019).

42. El-gebali, S. et al. The Pfam protein families database in 2019. 47, 427–432 (2019).

43. Guo, Y., Liu, C.-X., Zhang, L.-S., Wang, M.-Q. & Chen, H.-Y. Sterol content in the artificial diet of Mythimna separata affects the metabolomics of Arma chinensis (Fallou) as determined by proton nuclear magnetic resonance spectroscopy. Arch. Insect Biochem. Physiol. 96, (2017).

44. Ikekawa, N., Morisaki, M. & Fujimoto, Y. Sterol Metabolism in Insects: Dealkylation of Phytosterol to Cholesterol. Acc. Chem. Res. 26, 139–146 (1993).

45. Svoboda, J. A. & Weirich, G. F. Sterol metabolism in the tobacco hornworm, Manduca sexta--a review. Lipids 30, 263–7 (1995).

46. Avidor-Reiss, T. et al. Decoding cilia function: Defining specialized genes required for compartmentalized cilia biogenesis. Cell 117, 527–539 (2004).

47. Baughman, J. M. et al. Integrative genomics identifies MCU as an essential component of the mitochondrial calcium uniporter. Nature 476, 341–5 (2011).

48. Tsai, I. J. et al. The genomes of four tapeworm species reveal adaptations to parasitism. Nature 496, 57–63 (2013).

49. Berriman, M. et al. The genome of the blood fluke Schistosoma mansoni. 460, (2009).

50. Frayha, G. J. SHORT COMMUNICATION COMPARATIVE METABOLISM OF ACETATE IN THE TAENIID TAPEWORMS ECHINOCOCCUS GRANULOSUS,. 39, 167–170 (1971).

51. Žárský, V. & Tachezy, J. Evolutionary loss of peroxisomes - not limited to parasites. Biol. Direct 10, (2015).

52. Gabaldón, T., Ginger, M. L. & Michels, P. A. M. Molecular & Biochemical Parasitology Peroxisomes in parasitic protists. Mol. Biochem. Parasitol. 209, 35–45 (2016).

53. Castro, H. et al. Acta Tropica Functional insight into the glycosomal peroxiredoxin of Leishmania. Acta Trop. 201, 105217 (2020).

54. Dawidowski, M. et al. Inhibitors of PEX14 disrupt protein import into glycosomes and kill Trypanosoma parasites. 1420, 1416–1420 (2017).

55. Kalel, V. C. et al. BBA - Molecular Cell Research Evolutionary divergent PEX3 is essential for glycosome biogenesis and survival of trypanosomatid parasites. BBA - Mol. Cell Res. 1866, 118520 (2019).

56. Wu, B. et al. The heme biosynthetic pathway of the obligate Wolbachia endosymbiont of Brugia malayi as a potential anti-filarial drug target. PLoS Negl. Trop. Dis. 3, e475 (2009).

57. Goldberg, D. E. & Sigala, P. A. Plasmodium heme biosynthesis?: To be or not to be essential? 2–7 (2017).

58. Li, P., Xie, L., Gu, Y., Li, J. & Xie, J. Roles of Multifunctional COP9 Signalosome Complex in Cell Fate and Implications for Drug Discovery. 1246–1253 (2017). doi:10.1002/jcp.25696

59. Renatus, M. et al. Targeted inhibition of the COP9 signalosome for treatment of cancer. (2016). doi:10.1038/ncomms13166

60. Pereira, R. V., De Gomes, M. S., Jannotti-Passos, L. K., Borges, W. C. & Guerra-Sá, R. Characterisation of the COP9 signalosome in Schistosoma mansoni parasites. Parasitol. Res. 112, 2245–2253 (2013).

61. Hetfeld, B. K. J. et al. The COP9 signalosome-mediated deneddylation is stimulated by caspases during apoptosis. Apoptosis 13, 187–95 (2008).

62. Yuan, J. et al. RNA-dependent conversion of phosphoserine forms selenocysteine in eukaryotes and archaea. Proc. Natl. Acad. Sci. U. S. A. 103, 18923–18927 (2006).

63. Liang, P., Nair, J. R., Lei, S., McGuire, J. J. & Dolnick, B. J. Comparative genomic analysis reveals a novel mitochondrial isoform of human rTS protein and unusual phylogenetic distribution of the rTS gene. BMC Genomics 6, (2005).

64. Moricca, S. et al. Endemic and emerging pathogens threatening cork oak trees: Management options for conserving a unique forest ecosystem. Plant Dis. 100, 2184–2193 (2016).

65. Yin, Z. T. et al. Revisiting avian ‘missing’ genes from de novo assembled transcripts. BMC Genomics 20, (2019).

66. O’Leary, N. A. et al. Reference sequence (RefSeq) database at NCBI: Current status, taxonomic expansion, and functional annotation. Nucleic Acids Res. 44, D733–D745 (2016).

67. Camacho, C. et al. BLAST+: Architecture and applications. BMC Bioinformatics 10, (2009).

